# Bayesian Multi-Study Non-Negative Matrix Factorization for Mutational Signatures

**DOI:** 10.1101/2023.03.28.534619

**Authors:** Isabella N. Grabski, Lorenzo Trippa, Giovanni Parmigiani

## Abstract

Mutational signatures shed insight into the range of mutational processes giving rise to tumors and allow a better understanding of cancer origin. They are typically identified from high-throughput sequencing data of cancer genomes using non-negative matrix factorization (NMF), and many such techniques have been developed towards this aim. However, it is often of particular interest to compare mutational signatures across multiple conditions, e.g. to understand which signatures are present across different treatments, or to identify signatures that are shared or specific across cancer types. Existing techniques within the NMF context only allow decomposition within a single dataset, so that integrating results across multiple conditions requires running separate analyses on each dataset, followed by subjective and manual comparisons of the identified signatures. To address this issue, we propose a Bayesian multi-study NMF method that jointly decomposes multiple studies or conditions to identify signatures that are common, specific, or partially shared by any subset. We propose two models: a “discovery-only” model that estimates de novo signatures in a completely unsupervised manner, and a “recovery-discovery” model that builds informative priors from previously known signatures to both update the estimates of these signatures and identify any novel signatures. We then further extend these models to estimate the effects of sample-level covariates on the exposures to each signature, enforcing sparsity through a non-local spike-and-slab prior. We demonstrate our approach on a range of simulations, and apply our method to colorectal cancer samples to show its utility.

## 1 Introduction

The somatic mutations found in tumor genomes are the result of many mutational processes acting in concert. These processes can arise from a number of possible sources, including inaccuracies in DNA replication, DNA modification by enzymes, defective DNA repair, and exposure to mutagenic agents, such as tobacco smoking or ultraviolet light radiation [Alexandrov et al., 2013]. Identifying and characterizing these processes is an important step in understanding how various types of cancers arise. Moreover, the particular processes that contributed to any given individual’s cancer development could have key implications for prognosis and/or treatment.

Different processes have been shown to give rise to different patterns of mutations, which are termed mutational signatures. A mutational signature expresses the propensity of a particular process to produce various types of mutations and can be mathematically represented as a vector of rates for each type of mutation under consideration. Nik-Zainal et al. [2012] and Alexandrov et al. [2013] introduced the usage of non-negative matrix factorization (NMF) to estimate these signatures from sequenced tumor genomes. NMF decomposes mutation count data into a product of two lower-dimensional matrices: one represents the signatures and the other represents the exposures, i.e. the extent to which each signature contributed to each tumor’s accumulation of mutations.

Since then, a number of approaches using NMF have been developed and employed for signature estimation. Some, like the original approach of Nik-Zainal et al. [2012] and Alexandrov et al. [2013], use optimization techniques to identify the two lower-dimensional matrices [Alexandrov et al., 2020]. Recent developments in this area include an approach that explicitly accounts for background noise and uses regularization to enforce sparsity [Lal et al., 2021], and an approach that incorporates the sampling likelihood with regularization and informative priors to fine-tune model complexity [Li et al., 2020]. Fischer et al. [2013] introduced a probabilistic variant assuming the Poisson likelihood, and Rosales et al. [2017] generalized this Poisson model using an empirical Bayesian approach. Other Bayesian approaches include Kasar et al. [2015]. There are also some methods that estimate signatures with an entirely different framework, using topic models, e.g. through hierarchical Dirichlet processes [Shiraishi et al., 2015, Teh et al., 2006, Roberts, 2018], instead of NMF.

However, there has been very limited work on extending mutational signatures analysis to the multi-study context. This includes analyzing data from multiple studies, groups, or conditions to understand what signatures are shared or specific across these categories. For example, we may be interested in jointly analyzing breast cancer datasets from different populations, or datasets spanning multiple cancer types. Although grouping structure can be incorporated into models based on the hierarchical Dirichlet process [Teh et al., 2006, Roberts, 2018], existing NMF-based methods only consider a single data matrix. As such, analyzing multiple groups with NMF requires running separate analyses on each dataset and then subjectively comparing the identified signatures to determine what is shared and what is unique. Alternatively, multiple datasets can be concatenated into one and decomposed, but this again requires heuristic post-processing to understand how signatures are shared. These *ad hoc* solutions result in a loss of efficiency and fail to propagate important uncertainties from one step to the next.

In many cases, it is also of interest to compare the signatures estimated from one or more datasets to previously known signatures, such as those in the COSMIC database [Alexandrov et al., 2020]. This again generally requires heuristic approaches, such as considering an estimated signature the same as a previously known signature if the cosine similarity exceeds a pre-determined threshold. One extension of the hierarchical Dirichlet process allows “freezing” nodes that represent previously known signatures to effectively encode priors around these signatures [Roberts, 2018]. However, there is no existing equivalent based on NMF that both incorporates prior information about signatures and updates their estimates, allowing for variability around the provided values.

Finally, we may also often be interested in understanding the relationships between signatures and tumor-level covariates in one or more datasets. For example, we may want to know whether covariates such as the patient’s sex, smoking history, or the presence of homologous repair deficiency affect the expected exposure for a certain signature. Much existing work has correlated estimated exposures with patient features, but this does not allow the covariates to inform the matrix factorization, and does not propagate uncertainty. While the Tumor Covariate Signature Model [Robinson et al., 2019] does estimate the effects of covariates within a topic model, this has not been extended to the multi-study context. Recent work [Park et al., 2023] also estimates covariate effects for mutational signatures, but is also based in the single-study context and requires signatures to be specified ahead of time, rather than simultaneously discovering them.

Here, we introduce the first comprehensive multi-study NMF framework for mutational signatures estimation to address these gaps. In particular, we employ a Tetris-like prior [Grabski et al., 2020] to extend the Poisson model of Rosales et al. [2017] to the multi-study context by explicitly modeling the shared ownership of signatures. We propose two versions of this model: a “discovery-only” model in which all signatures are estimated *de novo* and a “recovery-discovery” model in which signatures are estimated both de novo and from informative priors built from existing signatures. Finally, we extend both of these models to incorporate tumor-level covariates, with a non-local prior based on the spike-and-slab prior of George and McCulloch [1993] to enforce sparsity in the coefficients. Because we simultaneously estimate signatures and covariate effects, this approach offers the opportunity for covariates to improve signature estimation. We compare these models to existing approaches on an extensive set of simulations, and demonstrate the utility of our method in colorectal cancer samples.

## 2 Results

### 2.1 A comprehensive multi-study NMF framework for mutational signatures estimation

We show a schematic in Figure 1 to illustrate our NMF framework. In the discovery-only model (Figure 1A), any number of counts matrices are taken as input, where each corresponds to a different study (or, equivalently, condition or group). Our method then jointly decomposes these studies under a Bayesian framework to estimate three key quantities: the set of signatures that appear in at least one study, a signature indicator matrix that identifies which signatures are present in each study, and the study-specific exposures for each sample in every study to each present signature. By estimating these quantities simultaneously, including an explicit representation of ownership over the signatures, this approach avoids having to run NMF on each study individually and compare results in an ad-hoc manner.

**Figure 1:**
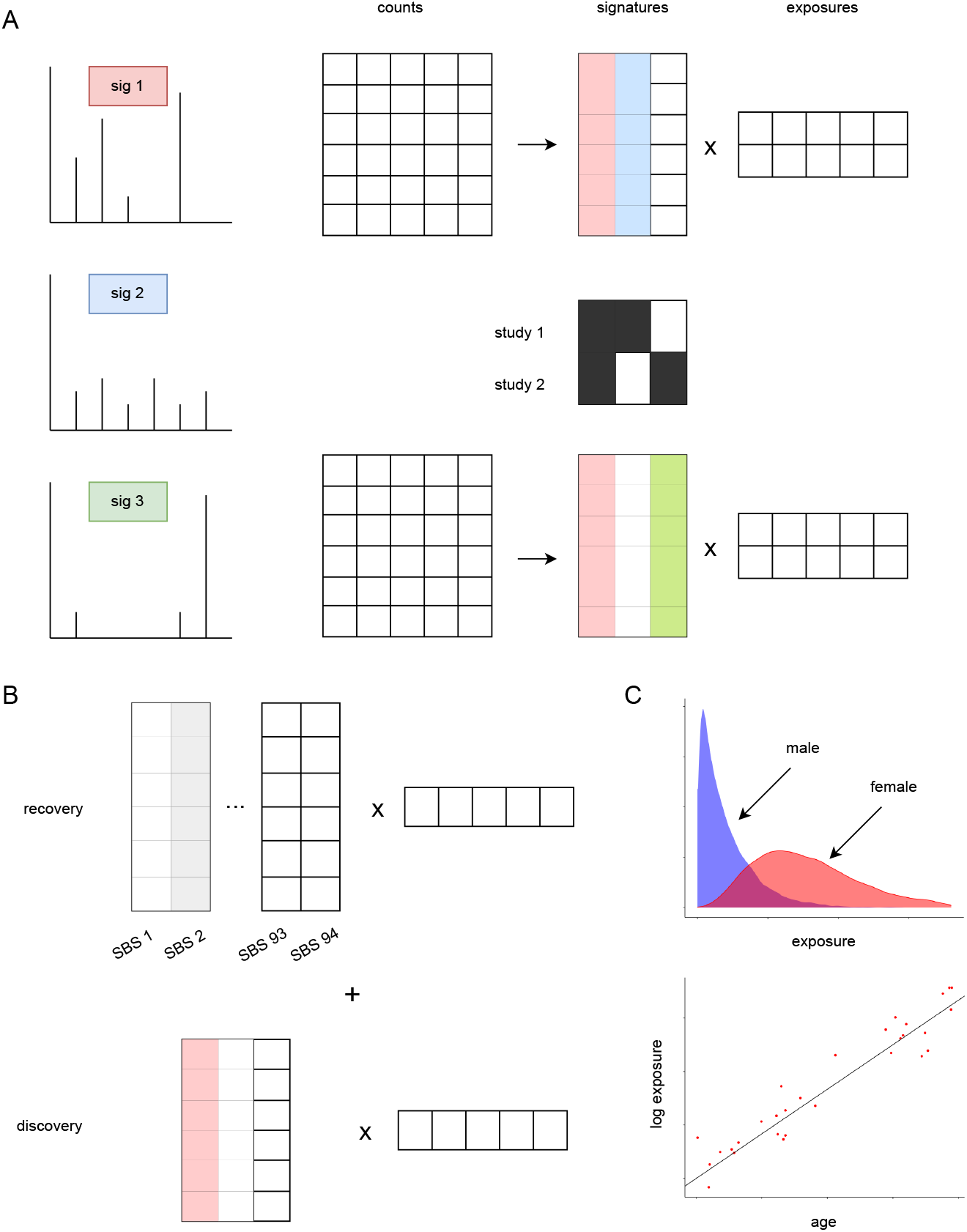
Schematic of our multi-study NMF framework. (A) Under the discovery-only model, we take the counts matrices for any number of studies as input, and learn signatures, exposures, and indicators of which signatures belong to each study. In this illustration, two studies are decomposed into a total of three signatures; the top study is found to contain the first two signatures, whereas the bottom study is found to contain the first and third signatures. This sharing pattern is shown by the signature indicator matrix. (B) Under the recovery-discovery model, we expand the NMF decomposition to both recover previously known signatures, encoded via informative priors, and to discover new signatures. Here, we show an example decomposition for one study, where one signature (SBS 2) is recovered and one signature is discovered. (C) In both models, we can incorporate covariates, such as gender and age as illustrated here, to learn relationships with the exposures.

In many settings, we are also interested in specifically determining whether any previously found signatures are present in the data. Under the recovery-discovery model (Figure 1B), we again take any number of counts matrices as input for joint estimation. However, we now expand the NMF decomposition into two terms, which we label as recovery and discovery respectively. The discovery term represents the de novo signatures, and is identical to the decomposition from the discovery-only model. The recovery term represents the previously found signatures and encodes the COSMIC v3 single-base substitution signatures [Alexandrov et al., 2020] through informative priors, unlike the mildly informative priors used in the discovery term. This encourages the estimation of signatures that are very similar to those in the COSMIC v3 database, while still allowing their values to be updated based on the provided new data. Hence, under this model, we learn signature estimates, signature indicator matrices, and exposures for both a set of discovered signatures and a set of recovered signatures.

Finally, we allow both models to take in optional, sample-level covariates and learn their associations with the exposures (Figure 1C). In particular, we model covariates as having multiplicative effects on the exposure distributions; since we use the Gamma distribution as the exposure prior, this can be interpreted as a multiplicative effect on the mean exposure. However, we choose a prior on the coefficients, specifically a non-local formulation of the spike-and-slab prior [George and McCulloch, 1993], that enforces sparsity in these effects. This improves interpretability as well as biological plausibility in the model by encouraging covariates to have non-zero effects on the exposures to a small set of signatures, rather than having potentially minor effects on the exposures to all signatures.

We provide further details on our models and inferential procedures in section 4.

### 2.2 Our approach improves signature estimation in the multi-study setting

To evaluate the performance of our approach first in the simpler setting without covariates, we developed a simulation study motivated by exploratory analyses of real data from different cancer types. In particular, we considered two main scenarios: a three-study simulation and a ten-study simulation. For each scenario, we generated 10 datasets and ran both our discovery-only and recovery-discovery methods, alongside existing methods. In particular, we compared our performance to two popular single-study approaches, signeR [Rosales et al., 2017] and sigProfiler [Alexandrov et al., 2020]; a multi-study approach, HDP [Roberts, 2018], that is outside of the NMF framework; and a semi-supervised version of HDP using frozen nodes to encode known signatures, which we refer to as HDP-frozen. When running the single-study methods, we carried out separate runs on each study. Further details on simulation set-up and running existing methods are in section 4.

We show results for both simulation scenarios in Figure 2 and examine three metrics. Namely, we report sensitivity (the proportion of ground-truth signatures for which sufficiently similar signatures were estimated in the correct study), precision (the proportion of estimated signatures that were sufficiently similar to ground-truth signatures in the correct study), and cosine similarity of estimated signatures with ground-truth signatures.

**Figure 2:**
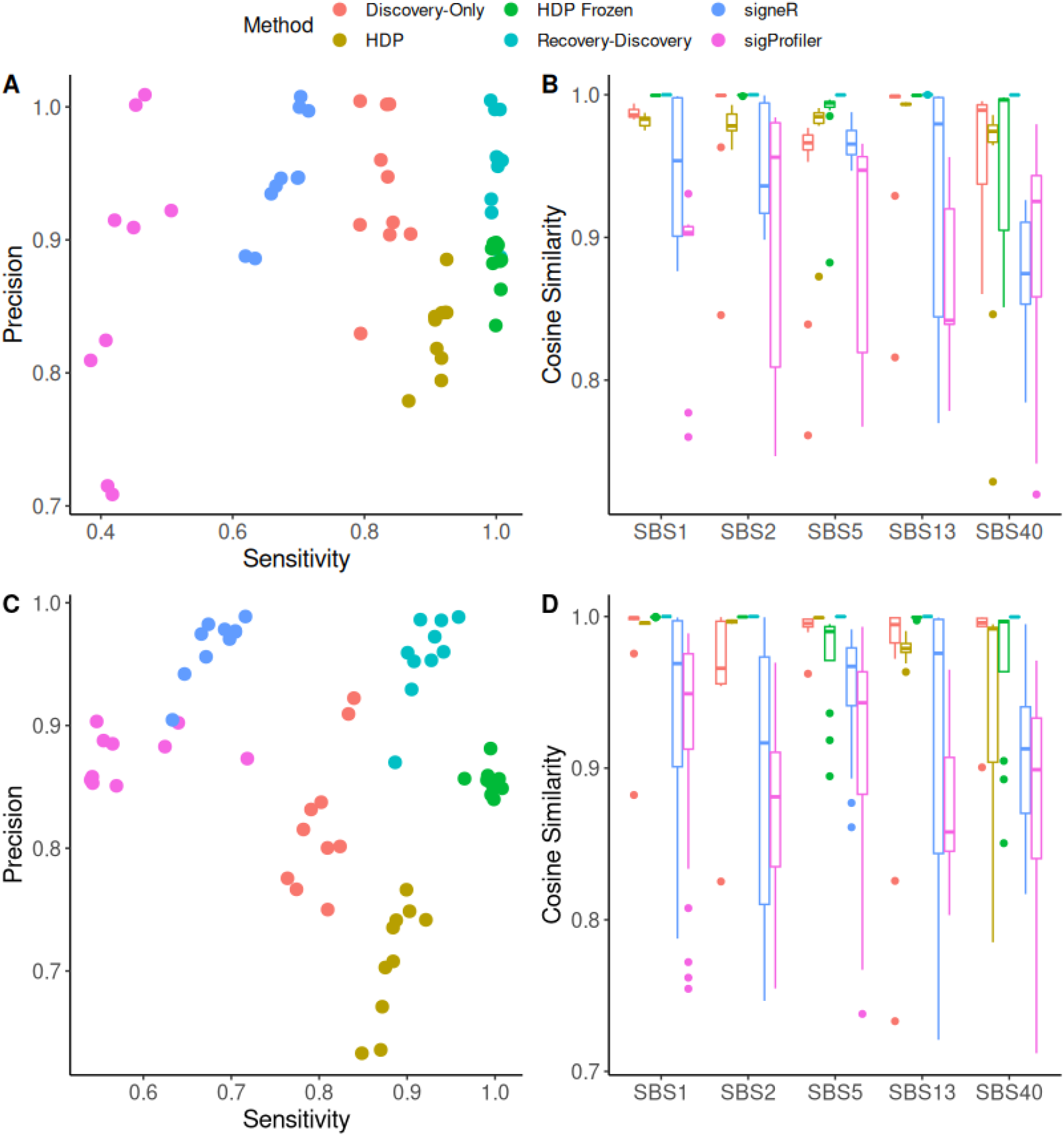
Simulation results in the setting without covariates. (A) Sensitivity and precision for the three-study simulation, comparing our discovery-only and recovery-discovery approaches to four existing methods: two single-study approaches (signeR and sigProfiler) and two multi-study approaches (HDP and HDP Frozen). Points are jittered for clarity. Only 9 points are shown for sigProfiler due to failure to converge in one simulation. (B) Cosine similarities in the three-study simulation for signatures that are estimated under all approaches. (C) Sensitivity and precision for the ten-study simulation. (D) Cosine similarities in the ten-study simulation for signatures that are estimated under all approaches.

In both the three-study and ten-study simulations, our discovery-only model had better sensitivity and precision than both single-study methods. This demonstrates the improvements of a multi-study approach in borrowing strength across datasets. When compared to the other fully unsupervised multi-study approach, HDP, the discovery-only model had overall similar performance under both simulations, representing a different trade-off between sensitivity and precision. Because this is a benchmark in the setting without covariates, the similar performance between the two is to be expected. Our recovery-discovery model had even better results than the discovery-only model, most notably in terms of sensitivity. Since the ground-truth signatures were taken from the COSMIC database, this model’s informative priors made it easier to capture weak signals such as SBS17a that were otherwise challenging to learn under any of the unsupervised approaches. While the results can again be interpreted as representing a different trade-off between sensitivity and precision in the ten-study simulation, under the three-study simulation, the recovery-discovery model showed comparable sensitivity but better precision than HDP-frozen.

Finally, when examining a set of signatures that were consistently found across all methods under consideration, our recovery-discovery model produced estimates with the highest cosine similarity to the ground-truth signatures across the board. This performance was closely followed by HDP-frozen. Among the fully unsupervised methods, our discovery-only model had the best estimates in four of the five signatures considered in the three-study simulation, and in three of the five signatures considered in the ten-study simulation. It is also notable that all of the multi-study approaches had much lower variance in cosine similarities than the single-study approaches, showing that leveraging multiple studies improves both accuracy and precision.

We also conducted a simulation study to further characterize the properties of the informative priors used in the recovery-discovery model and to better understand their performance. In particular, for each of the 78 signatures in the COSMIC database, we generated five single-study datasets where all signal comes from a perturbed version of that signature. For example, this could arise in a setting with batch effects. We then ran the recovery-discovery model on each of these single-study datasets.

The results are shown in Figure 3. In Figure 3A, we highlight three patterns. First, in many cases, the cosine similarity of the estimated signature with the reference signature was very close to 1, even when the data-generating signature had a much lower cosine similarity with the reference signature. This indicates that the informative prior had a strong enough influence as to shrink the estimate to the reference, despite the noise in the data-generating process. In these cases, in addition to the signature in the recovery term, a second signature was also found in the discovery term. These discovered signatures were nearly always much less similar to the reference signature as the recovered signatures (Figure 3B). This suggests that these discovered signatures were fitting to the extraneous noise from the data-generating process, possibly sharpening the estimation of the recovered signatures.

**Figure 3:**
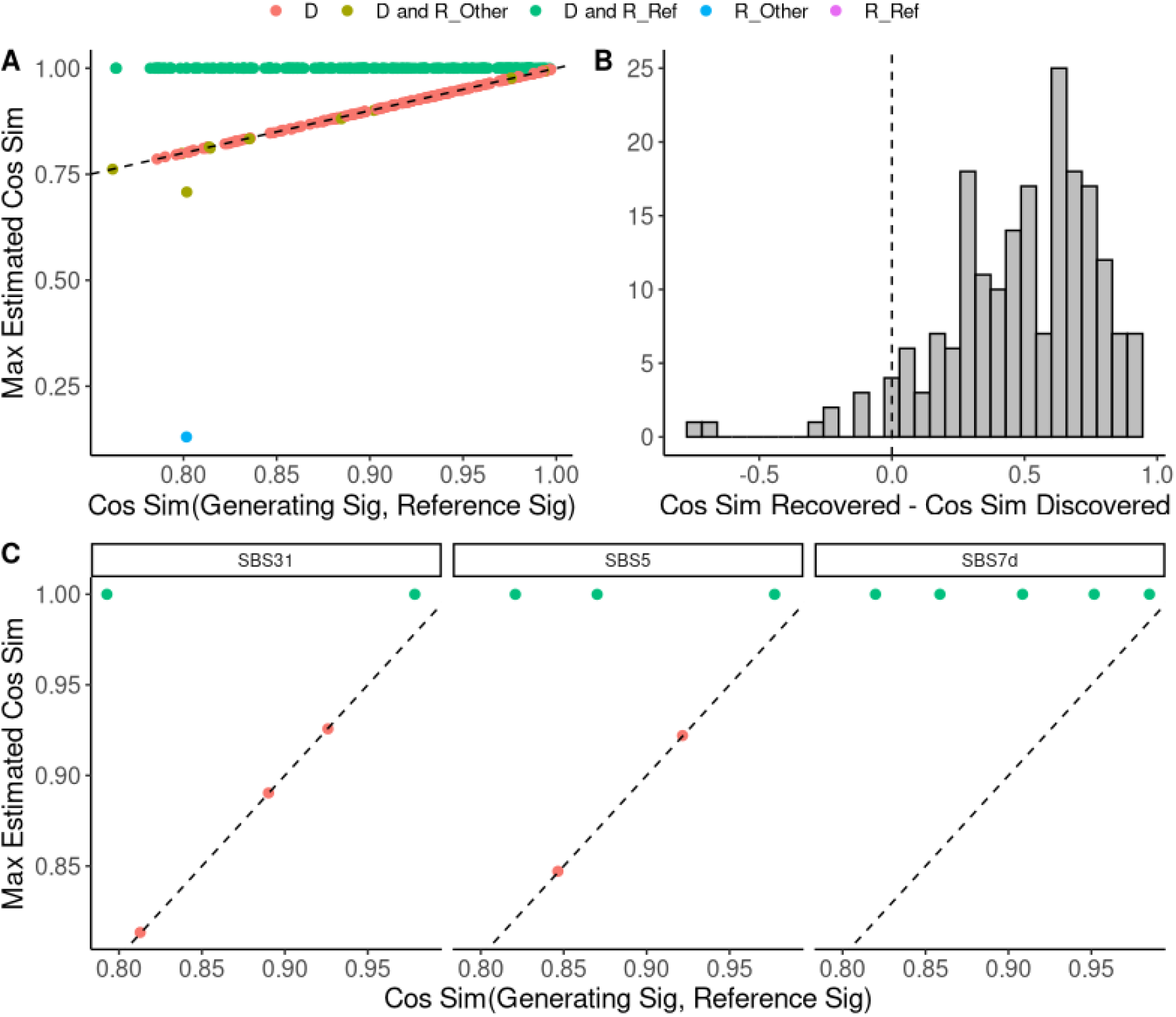
Simulation results from running the recovery-discovery model on single-study data generated from perturbed versions of the reference signatures. (A) Comparison of the cosine similarity between the data-generating signature and the reference signature, indicating the extent of perturbation, to the cosine similarity between the estimated signature under our model and the reference signature. If more than one signature was found in a single dataset, we report the maximum of all cosine similarities, such that one and only one point per dataset is included. Colors indicate whether the estimated signature was in the discovery term (“D”), in the recovery term in the slot corresponding to the reference signature (“R_Ref”), in the recovery term in a slot corresponding to a different signature (“R_Other”), or signatures were estimated in both the discovery and recovery terms (“D and R_Ref” and “D and R_Other”). (B) Difference in cosine similarity to the reference signature of the signature in the recovery term and the signature in the discovery term, for cases in which a signature was found in both the discovery and recovery terms (i.e. points indicated as “D and R_Ref” or “D and R_Other” in panel A). (C) Comparison of the cosine similarity between the data-generating signature and the reference signature, to the cosine similarity of the estimated signature under our model and the reference signature, for the five datasets corresponding to three specific signatures.

Second, in many other cases, the cosine similarity of the estimated signature with the reference signature was very close to that between the data-generating signature and the reference signature. This suggests that our approach captured the data-generating signature, rather than performing shrinkage to the reference. In the majority of these cases, a signature was captured in the discovery term with no signatures in the recovery term. In some instances, a signature was found in the recovery term in a slot corresponding to another signature, possibly with an additional signature in the discovery term. Finally, in just one case, the cosine similarity of the estimated signature with the reference signature was very low, indicating a poor fit. In this case, a signature was found in the recovery term, in a slot corresponding to a different signature from the reference. This suggests inadvertent shrinkage to a different reference signature.

Figure 3C shows the results for three signatures, as specific examples. The implied strength of the informative prior appears to vary across signatures. For example, for SBS7d, a near-perfect estimate is found in every case regardless of the size of perturbation. By contrast, SBS5 and SBS31 have variable results across the range of tested perturbations, suggesting that some degree of noise may influence the outcome. Collectively, these results help characterize when and how the recovery-discovery model recovers known signatures in ambiguous scenarios, and shed insight into what estimation might reveal in real data settings where, hypothetically, a cancer- or tissue-specific version (i.e. a perturbation) of a signature may be present.

### 2.3 Our approach accurately detects and estimates covariate effects

In the setting with covariates, we conducted a simulation study to assess how well our model identifies which covariates have non-zero effects on which signatures, as well as how accurately the corresponding coefficients are estimated. This simulation study was motivated by the colorectal cancer application, thus matching the dimensionality of those three studies and containing signatures and covariate effects likely to be present in each. Further details are in section 4.

First, we describe estimation of the signatures and sharing pattern. In the discovery-only model, out of ten runs of simulations using the same data-generating set-up, the ground-truth set of signatures with exactly the correct sharing pattern across the three studies was found in half of the runs. The other runs found a sparser solution, in which the signature SBS10b was omitted but the other signatures were still estimated with the correct sharing pattern. We ran the recovery-discovery model on the same data, and found the correct solution in all runs. The greater sensitivity to SBS10b, which was difficult to detect, can be attributed to the informative priors of the recovery-discovery model.

Next, we examine identification and estimation of covariate effects. In this simulation, non-null effects from the covariates (gender and age) were only present in one of the studies. Because this simulation is modeled after the real data application, the study with non-null effects only has 16 samples, which offers an opportunity to demonstrate our approach’s utility in the small-sample setting. Under both the discovery-only and recovery-discovery models, covariate effects in all signatures for the other two studies had posterior inclusion probabilities (PIPs) under 0.1%, correctly indicating with low uncertainty that there are no effects in that study. Results for both the discovery-only and the recovery-discovery model in the study that does have non-null effects are shown in Figure 4. Both models found PIPs close to 1 for signatures with ground-truth non-null effects, and PIPs close to 0 for ground-truth null effects. This again shows that our sparsity-encouraging prior enables our approach to distinguish non-null and null signal effectively. We also show that both models accurately estimate the coefficient estimates in all cases. It should be noted that the estimation of the gender effect for SBS10b tended to be more variable both in the posterior inclusion probability and in the coefficient. This can be attributed to the fact that the exposures to this signature were on average smaller than those to SBS10a, which makes estimation more challenging. However, this variability was lower under the recovery-discovery model, suggesting that the semi-supervised inference of the signatures also lent more stability to estimating the covariate effects.

**Figure 4:**
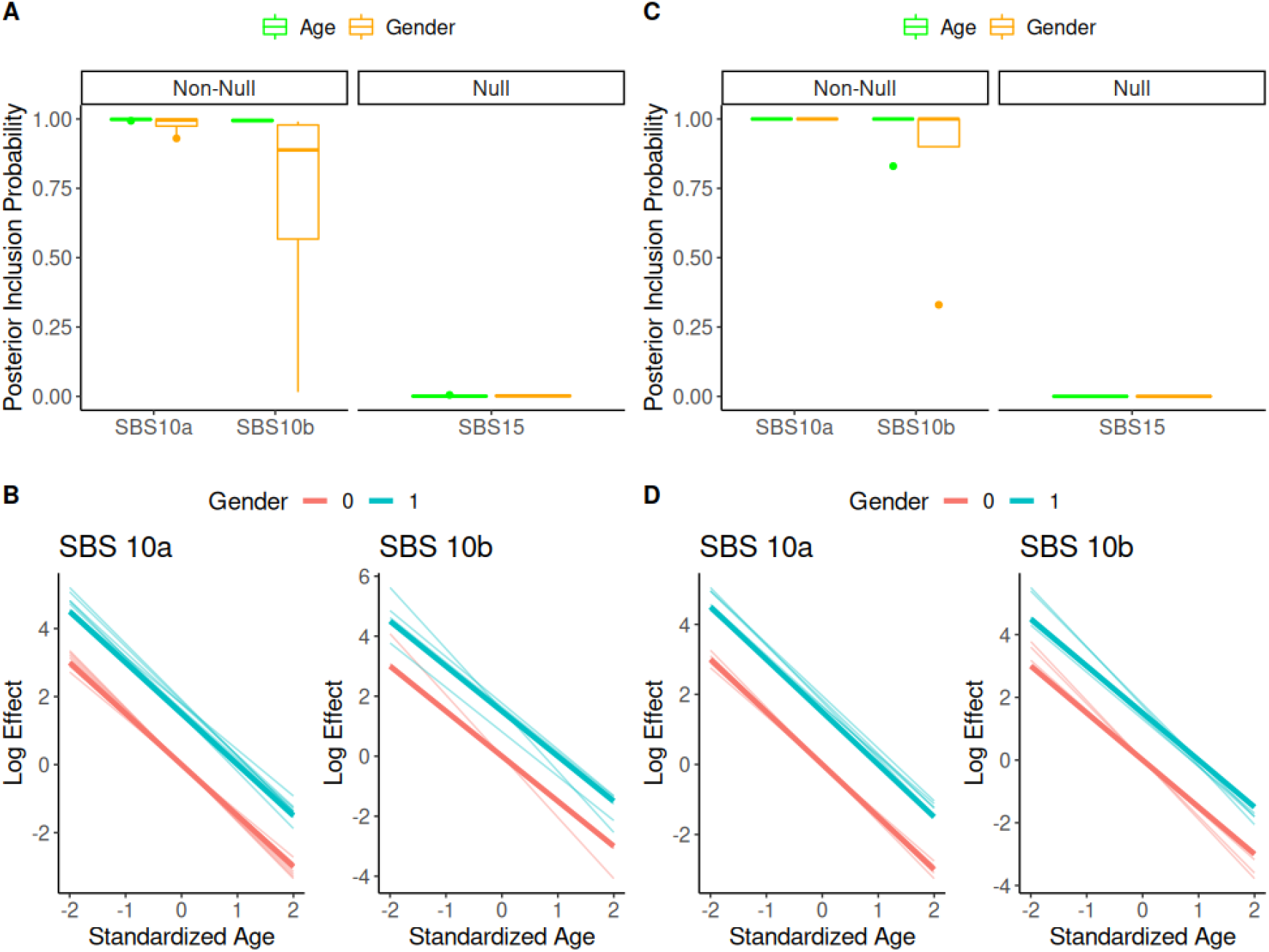
Results from a simulation with covariates, using 10 simulated datasets. (A) Under the discovery-only model, posterior inclusion probabilities for the two covariates (gender and age) in a study with ground-truth non-null effects in two signatures and a ground-truth null effect in the other signature. (B) Under the discovery-only model, estimated associations with gender and age for the two signatures with non-null effects. The true associations are indicated by the dark blue and dark red lines, and the lighter opacity lines indicate estimated associations from each simulation run. (C) Under the recovery-discovery model, posterior inclusion probabilities. (D) Under the recovery-discovery model, estimated associations.

### 2.4 Our approach reveals shared and distinct mechanisms in colorectal cancer

We demonstrate our approach on real data by applying it to a set of important questions in colorectal cancer (CRC). Many different mutational signatures have been previously identified in CRC samples, from those associated with aging to those associated with the failure of endogenous molecular processes [Díaz-Gay and Alexandrov, 2021]. In particular, two distinct classes of CRC samples, characterized by their high mutational burden as “hypermutated” and “ultra-hypermutated” respectively, have been found to contain specific mutational processes. Hypermutated samples are associated with microsatellite instability (MSI), and ultra-hypermutated samples are associated with POLE (polymerase epsilon) mutations, which affect proofreading in DNA replication Díaz-Gay and Alexandrov [2021]. Despite the characterization of these processes, much remains unknown about their relationship with epidemiological factors. For example, while POLE mutations have been associated with younger age and male sex [Puccini et al., 2017], there has been limited work examining relationships between covariates such as these and the repertoire of mutational signatures in CRC.

To address questions like these, we applied our recovery-discovery model to 406 CRC samples from the TCGA Research Network. Specifically, our goals were to precisely identify which signatures are shared and distinct across the mutational burden classes, and to understand what relationships, if any, exist with gender and age. Our approach offers a unique opportunity to investigate these questions because of our explicit representation of the signatures sharing pattern across classes, as well as our ability to simultaneously estimate covariate effects in a way that can influence signature estimation too. We divided the samples into three “studies” or groups based on their tumor mutational burden: non-hypermutated (328 samples), hypermutated (62 samples), and ultra-hypermutated (16 samples), following guidance from Díaz-Gay and Alexandrov [2021]. As covariates, we included gender coded as a binary variable and age coded as a z-scored continuous variable (denoted hereafter as age-continuous). Because initial exploratory analyses indicated that some signatures may have a changing relationship with age in the oldest patients, we also included a binary indicator of whether a patient is older than 75 (denoted as age-step), and an interaction term between age-step and age-continuous (denoted as age-interaction).

Results from the recovery-discovery model are shown in Figure 5A. Our model found a total of 23 signatures, two in the discovery component and 21 in the recovery component. The “discovered” signatures were labeled by their closest match in cosine similarity to the COSMIC v3 signatures, as long as the cosine similarity was over 0.70. Many of the signatures found can be divided into 3 main categories: clock-like signatures (SBS 1, SBS 5, SBS 40), MSI (SBS 6, SBS 14, SBS 15, SBS 20, SBS 21), and POLE mutations (SBS 10a, SBS 10b, SBS 28), all of which are expected in CRC based on previous work [Díaz-Gay and Alexandrov, 2021].

**Figure 5:**
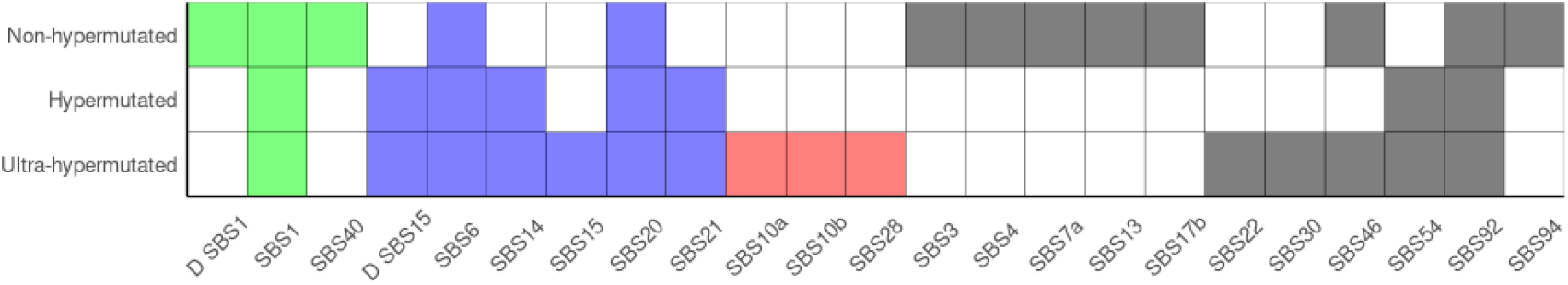
Signature indicator matrix estimated using the recovery-discovery model in the colorectal cancer application. Green indicates clock-like signatures, red indicates MSI signatures, blue indicates POLE mutation signatures, and gray indicates signatures falling into other categories. A “D” at the front of the signature name indicates that it was found in the discovery component.

The sharing pattern of signatures in each of these categories sheds insight into the presence of these mutational processes across the three groups. First, clock-like signatures are found in all three groups, but with some variation. SBS 1 in the recovery component was shared by all three groups, but both SBS 40 in the recovery component and SBS 1 in the discovery component were specific to the non-hypermutated group. This suggests that while aging plays a role across the board, there are specific age-related mechanisms in the non-hypermutated samples. In particular, the repetition of SBS 1 in both the discovery and recovery components, in light of the previously discussed simulation results with perturbed signatures, may indicate the presence of a non-hypermutated-specific version of this signature.

Next, most of the MSI signatures are not found in the non-hypermutated samples. This finding is consistent with previous reports, which implicate MSI as a feature of hypermutation [Díaz-Gay and Alexandrov, 2021]. However, our result adds greater nuance to this by specifically finding that while five of the six MSI signatures are shared by both the ultra-hypermutated and the hypermutated groups, SBS 15 in the recovery component is found only in the ultra-hypermutated group. This suggests that there may be at least one MSI-related mechanism that only contributes to ultra-hypermutation. It should be noted that SBS 15 in the discovery component was shared by both groups, which could imply the existence of context-specific versions of this signature: one shared by both, and one only in the ultra-hypermutated. Finally, all of the POLE mutation signatures were only found in the ultra-hypermutated group, consistent with previous reports [Díaz-Gay and Alexandrov, 2021].

There were additional signatures recovered that don’t fall into any of these three categories. Namely, the nonhypermutated group was found to contain a DNA damage repair signature SBS 3, tobacco smoking signature SBS 4, ultraviolet radiation signature SBS 7a, APOBEC signature SBS 13, and two signatures with unknown etiology, SBS 17b and SBS 94. Several of these have been previously reported in cRC [Díaz-Gay and Alexandrov, 2021], often in small subsets of samples, but SBS 4 and SBS 7a are not traditionally associated with this cancer type. The ultra-hypermutated group was also found to contain an aristolochic acid exposure signature SBS 22 and a DNA repair signature SBS 30, the latter of which has been previously reported in CRC. Exposures to both of these signatures were very low (less than 2% of total exposure) in all but one sample in each case. This suggests that both are relatively rare mechanisms potentially exhibited by just one patient each. Finally, known sequencing artifacts SBS 46 and SBS 54 were each found in two of the groups, and another tobacco smoking signature SBS 92, which like SBS 4 is not traditionally associated with CRC, was found in all three groups. All together, these additional signatures across the three groups point to a heterogeneous range of mechanisms that may contribute to various subsets of patients, even though they are not among the driving forces traditionally ascribed to CRC. While many of these have been previously reported in this cancer type, others are potentially new, rare mechanisms. It should, however, be noted that these signatures without traditional associations could also possibly be unknown signatures that are simply similar enough to a signature in the COSMIC catalogue that they were pulled by the informative priors.

We summarize the effects of the covariates on exposures to all signatures in Figure 6. In the hypermutated group, the PIPs for all covariates on all signatures were less than 4%, indicating that there is most likely no relationship with any of our gender and age variables. However, both the ultra-hypermutated and the non-hypermutated groups had evidence of relationships.

**Figure 6:**
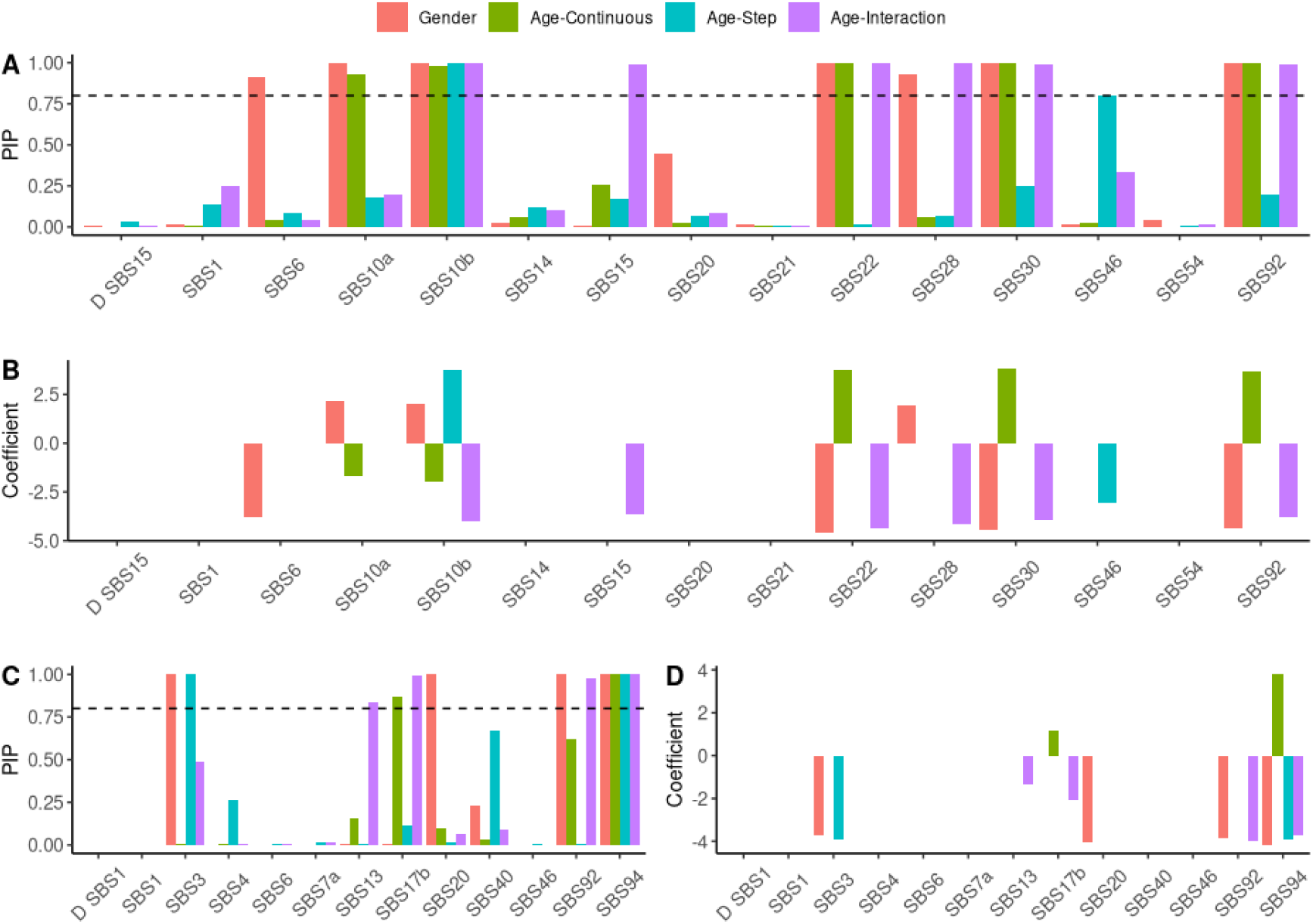
Effects of covariates on exposures to signatures in the ultra-hypermutated and non-hypermutated samples. (A) Posterior inclusion probabilities (PIPs) in the ultra-hypermutated group. (B) Coefficients in the ultra-hypermutated group for all cases where the PIP is greater than 0.8. (C) PIPs in the non-hypermutated group. (D) Coefficients in the non-hypermutated group for all cases where the PIP is greater than 0.8.

In the ultra-hypermutated group, most of the PIPs were either quite high (above 80%) or quite small (below 20%), reflecting the sparsity imposed by the nonlocal prior. Particularly high PIPs were seen in two of the MSI signatures, SBS 6 and SBS 15. Specifically, SBS 6 had a high PIP for gender, with a posterior mean coefficient of −3.8, suggesting a substantially lower exposure in males than females. SBS 15 had a high PIP for the age-interaction term, with a posterior mean coefficient of −3.7, which implies a decreasing relationship with age only among those older than 75. There were also high PIPs for the POLE mutation signatures. All three of SBS 10a, SBS 10b, and SBS 28 had high PIPs for gender, with posterior mean coefficients of 2.1, 2.0, and 1.9 respectively. Consistent with previous findings [Puccini et al., 2017], this suggests that male patients have larger exposures to these signatures. Each of the three also have high PIPs for one or more of the age-related covariates, with coefficient estimates implying decreasing exposures at older ages. This is consistent as well with previous findings that POLE mutations are less common among older patients [Puccini et al., 2017]. In the latter two signatures, the large and negative coefficients on the age-interaction terms specifically imply an even greater decreasing association with age among those older than 75. These varying associations even among related signatures suggest heterogeneity in how these mutational processes manifest in different demographics.

Finally, high PIPs were found in four of the other signatures that did not fall in the clock-like, MSI, or POLE mutation categories, namely SBS 22, SBS 30, SBS 46, and SBS 92. However, all of these signatures only had high exposures in a handful or even just one sample. Hence, these associations are driven by these limited observations are likely not to be generalizable.

In the non-hypermutated group, SBS 3, SBS 13, SBS 17b, SBS 92, and SBS 94 all had high PIPs. Exposure to SBS 3 is negatively associated with both gender and age, suggesting that this signature is more likely to be observed in females and younger patients. Exposure to SBS 94 is also more likely in female patients, but with a more complex association with age. Namely, there is an increasing association with age only among patients younger than 75. Because the remaining signatures SBS 13, SBS 17b, and SBS 92 have low exposures in the majority of patients, their associations may not be generalizable.

## 3 Discussion

We presented a comprehensive Bayesian multi-study NMF framework for mutational signatures to address three key challenges: extending NMF to the multi-study setting, implementing semi-supervised inference, and simultaneously estimating covariate effects. We showed the strengths of our approach against competing methods through a set of simulations. Finally, we demonstrated practical utility with a thorough analysis of colorectal cancer samples, in which we both recapitulated previously known biology and proposed new insights. This application particularly highlighted our approach’s ability to rigorously identify sharing patterns and associations that would have required ad-hoc analyses, and would have lacked systematic uncertainty assessment, using existing methods.

While our proposal represents a powerful and comprehensive approach to mutational signatures estimation, there are still some limitations. One key challenge lies in the multi-modality of the posterior. A naively implemented sampler can easily become stuck in a local posterior mode because of the high energy barrier required to move from one mode to another. In our approach, we employ tempering to more easily explore the parameter space, as well as marginal likelihood computations to probabilistically summarize alternative solutions. However, this introduces a substantial additional computational burden, which limits the applicability of our approach in very large settings. Alternative approaches to characterize the multi-modal posterior could be very valuable.

There are also several opportunities for extensions of our modeling approach. For example, our signature sharing matrix imposes regularization through the binary elements of *A.* This can be highly beneficial for interpretability, since it implies a signature is either present in a study or it’s not, but can over-simplify sharing patterns in scenarios where a more nuanced relationship is present. Hence, our approach offers substantial benefits in settings where binary sharing patterns are likely, but may be less appropriate if patterns across groups are more continuous in nature. Conversely, we do not enforce any sparsity in the exposures of individual samples to each signature, conditional on the signatures present. While this carries some computational advantages, it can be difficult to interpret which signatures meaningfully contribute to each sample within a study.

## 4 Methods

### 4.1 NMF for Mutational Signatures

In this work, we focus primarily on single-base substitution (SBS) mutational signatures. SBSs are mutations in which one nucleotide base is substituted for another. There are six possible such substitutions, namely C>A, C>G, C>T, T>A, T>C and T>G, where, for example, C>A denotes a C mutated into an A. However, the properties of these substitutions are often also affected by the bases on either side of the mutated site. Since there are four possible bases that could be on each side, we typically consider a total of 4 · 6 · 4 = 96 total mutational motifs [Alexandrov et al., 2013]. Note that it is possible to consider other types of mutational signatures, such as by increasing the number of bases considered on each side, or by instead examining double-base substitutions or indels. Although we ground the context of this work in SBS signatures for expositional simplicity, our methods are described and implemented more generally and can be applied with any classification of mutational motifs.

Mathematically, each mutational signature is represented as a *K* × 1 vector of rates, where *K* is the number of mutational motifs. The key assumption in the NMF approach is that the mutations in each tumor sample arise from the additive effects of the *N* mutational signatures at play. In particular, suppose we have a K × *G* matrix of mutational counts ***M***, where the (*k, g*)th entry indicates how many mutations of motif *k* were observed in sample *g*, for *k* = 1,…, *K* and *g* = 1,…, *G*. Then, under the NMF model, we can decompose

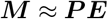

for *K* × *N* signatures matrix ***P*** and *N* × *G* exposures matrix ***E***. Here, each column of ***P*** is a *K* × 1 mutational signature, and the (*n*, *g*)th entry of ***E*** quantifies how much signature *n* contributed to the mutations of sample *g*.

Under the classical NMF approaches to this problem, Nik-Zainal et al. [2012] and Alexandrov et al. [2013] solved the optimization problem

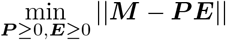

for chosen *N* and norm. However, it can be shown for a specific choice of norm, this optimization is equivalent to maximum likelihood estimation of ***P***, ***E*** under the model

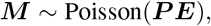

which was leveraged by Fischer et al. [2013] using an EM algorithm. Rosales et al. [2017] then introduced an empirical Bayesian treatment to this model, signeR, by combining MCMC and EM techniques. This approach results in improved uncertainty quantification compared to previous methods by estimating the posterior distributions for ***P*** and ***E***.

Although NMF methods have been proven to provide powerful and interpretable results, these existing approaches still have several important limitations: (1) they can only decompose a single data matrix at a time, (2) they cannot simultaneously estimate the presence of previously known and *de novo* signatures in a way that also updates the known signatures’ estimates, and (3) they cannot incorporate tumor-level covariates into the matrix factorization. Hence, we propose a comprehensive NMF framework that addresses all of these limitations via two multi-study models, which we call the discovery-only model and the recovery-discovery model, that can both be extended to incorporate covariate effects.

### 4.2 Discovery-Only Model

Under the discovery-only model, *S* datasets are jointly decomposed in an entirely unsupervised manner to estimate *de novo* signatures. In particular, if ***M**_s_* represents the *K* × *G_s_* matrix of mutational counts for the *G_s_* samples in study *s*, then we build on the Poisson model of Fischer et al. [2013] to assume

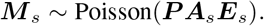

Here, ***P*** is the *K* × *N* matrix of signatures as defined in the previous subsection, where *N* is the total number of signatures present across all studies under analysis, and ***E**_s_* is the *N* × *G_s_* matrix of exposures for study *s*. Note that ***P*** is a common parameter across all studies.

The *N* × *N* study-level indicator matrix ***A**_s_* is our key element to estimate the pattern of sharing of signatures, as introduced in the context of factor analysis by Grabski et al. [2020]. This matrix consists of all 0s, except for the diagonal entries, which are either 1 or 0. The *n*th diagonal entry of ***A**_s_* is 1 whenever the *n*th signature is present in study *s*, so the product ***PA**_s_**E**_s_* will include the corresponding elements. Hence, ***A**_s_* controls which signatures are included and which are not into the model for study *s*.

We denote by 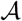 the overall *S* × *N* signature indicator matrix whose sth row consists of the *N* diagonal entries of ***A**_s_*. For example, if the *n*th column of 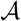 consists of all 1s, the *n*th signature is shared by all; if it consists of exactly a single 1, this signature only belongs to one study; and if it consists of more than one 1 and at least one 0, this signature is shared by some but not all studies.

Our prior model builds on signeR [Rosales et al., 2017]. In particular, we assume conjugate Gamma priors for the entries of ***P*** and ***E**_s_*, such that

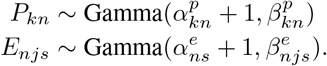

We then further assume hyperpriors for these shape and rate parameters, specifically

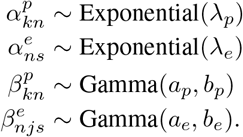

Although the Indian Buffet Process (IBP) was used as the prior on 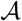 in the original factor analysis context [Grabski et al., 2020], we found in practice that mixing was more challenging with the IBP in this model. Instead, we use the related, but simpler, Beta-Bernoulli prior on each column of 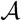, i.e.

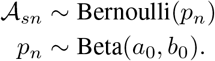

We fix the number of columns (i.e. the number of signatures) *N* to some large value, for example 50, but choose *a*_0_, *b*_0_ to encourage sparsity in 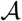. If all entries in a given column are 0, the corresponding signature can be interpreted as not present or contributing to any of the studies under consideration. As a result, the expected number of signatures is typically much smaller than the dimension *N*, and estimating 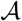 in this way allows us to automatically learn a posterior distribution over the total number of signatures that should be present. This is a substantial advantage over previous approaches, including signeR, in which the number of signatures is commonly selected by factorizing the data with different ranks and using a criterion such as BIC to select the best value.

In signeR, hyperparameter values are estimated from the data through an empirical Bayesian approach. While a similar approach could in principle be applied here, we instead found it sufficient to use default hyperparameter values of *a_p_* = 5, *b_p_* = 0.05, *a_e_* = 5, *b_e_* = 0.01, λ_*p*_ = 0.5, λ_*e*_ = 1, *a*_0_ = 0.8, and *b*_0_ = 0.8. Because different studies can potentially have highly varying total numbers of mutations, we add further stabilization by fixing a study-specific normalizing constant ***W**_s_* in the decomposition, i.e. as

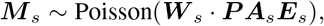

where · here indicates element-wise multiplication and ***W**_s_* is a *K* × *G_s_* matrix whose elements are all set to the median number of total mutations across all samples in the study. This can effectively be interpreted as normalizing exposures across studies, but because Gamma distributions form scale families, the implied priors on the exposures still follow Gamma distributions.

To estimate these parameters, we use a Metropolis-within-Gibbs sampler extended from the sampler used in signeR [Rosales et al., 2017]. Further details on the sampling steps can be found in the supplementary materials. To improve mixing, we use a tempering scheme in which the likelihood terms for the sampling step for 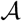 are raised to a power *γ* that gradually increases from 0 to 1 over the initial iterations. When *γ* = 0, this corresponds to sampling under the prior, and when *γ* = 1, this corresponds to sampling under the posterior. Our specific tempering scheme is as follows. The first 10 iterations are run with *γ* = 0, i.e. fully under the prior. Next, *γ* is set to 10^−*x*^ for *x* = 9, …, 5 for 100 iterations each. Then, *γ* = 10^−4^ for the next 400 iterations. Finally, *γ* is set to (1 + *x*) · 10^−*y*^ for *x* = 0, 0.1, …, 8.9 and *y* = 4, …, 1 for 100 iterations each. After this, *γ* = 1 for all remaining iterations. We found this scheme to encourage greater exploration of the parameter space and improve convergence.

Because the number of signatures as well as the sharing pattern can change from iteration to iteration, summarizing the posterior output is not straightforward. We choose point estimates to report by taking the five MCMC samples of 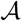 that occurred most often in the chain and running the sampler conditional on each of these values of 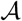. We estimate the marginal likelihood in each case by computing the harmonic mean estimator with parameter reduction [Raftery et al., 2006] (marginalizing out ***P***, ***E*** from the likelihood calculation in each iteration) from the conditional sampler output. We report the value of 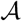 yielding the largest marginal likelihood, and summarize the remaining parameters as their posterior means from the conditional sampler output.

### 4.3 Recovery-Discovery Model

Under the recovery-discovery model, we jointly decompose *S* datasets to learn the presence of both *de novo* (discovered) and previously known (recovered) signatures. We implement this model using the known signatures from the COSMIC database [Alexandrov et al., 2020], but different or additional signatures can be easily incorporated as well. The recovery-discovery model expands the previously described model as

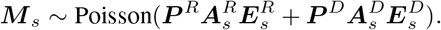

Here, the superscript *R* refers to parameters associated with the recovered (previously known) signatures, while the superscript *D* refers to parameters associated with the discovered (de novo) signatures.

The priors on ***P**^D^*, 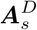, and 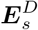 are exactly as described in the previous section. We also use the same priors for 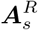 and 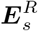. However, we instead build informative Gamma priors for each of the recovered signatures in the matrix ***P**^R^*. In particular, for each signature that we wish to include, we simulate a 500-sample dataset under our model using just the published estimate of that signature and a constant exposure of 500. Hence, this simulated dataset represents mutations that arise only from that signature. We then run the discovery-only sampler on this dataset with the number of signatures fixed to 1, and use the resulting learned posterior distribution of the signature estimate to set the shape and rate parameters of its prior.

With these priors in place, the MCMC sampler for this model is a simple extension of the one developed for the discovery-only case. We use the same hyperparameters for all terms associated with the discovery component, and the hyperparameters for the recovery parameters match those used for the corresponding discovery parameters. The exception is that to encourage the prioritization of recovered signatures over discovered signatures, we set the hyperparameters for 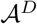 as 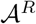 and the hyperparameters for 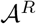 as 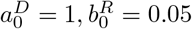. Detailed steps are in the supplementary materials.

### 4.4 Extensions with Covariates

We extended both the discovery-only and the recovery-discovery models to incorporate sample-level covariates. Suppose we have covariate vector ***x**_js_* for sample *j* in study *s.* Then we modify the exposure prior as

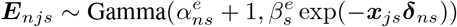

in the discovery-only case

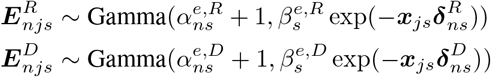

and in the recovery-discovery case.

Here, ***δ**_ns_* represents the vector of coefficients for the effects of each covariate on the exposure to signature *n* in study *s*. Nonzero values of ***δ**_ns_* can be interpreted as indicating multiplicative effects on the mean exposure. For example, if we had a single binary covariate *x_js_*, then a sample with *x_js_* = 1 has exp(*δ_ns_*) times the mean exposure to signature *n* as a sample with *x_js_* = 0.

It is important to enforce sparsity in the coefficients ***δ**_ns_*, since we expect most covariates to have effects on the exposures of particular, but not all, signatures. To do so, we use a product-moment prior version [Avalos-Pacheco et al., 2022] of the spike-and-slab prior from George and McCulloch [1993]. In particular, suppose the coefficient *δ_ins_* for covariate *i* arises from one of two possible distributions: a “spike” component, which is a low-variance normal distribution peaked sharply about 0, and a “slab” component, which is a higher variance normal distribution also centered at 0. We define indicator variables *γ_ins_* that express which component *δ_ins_* belongs to, which allows us to specify the prior

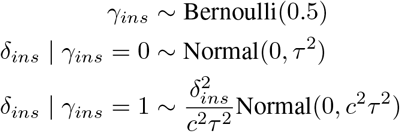

for appropriately chosen *c* and *τ*. Note that without the 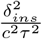 term, this would match the specification of George and McCulloch [1993].

As a default, we set *c* = 10 and *τ* = 0.3. To sample the coefficients, we leverage the results of Rossell and Telesca [2017], which shows that nonlocal priors can be represented as mixtures of truncated normals, giving rise to a simple sampling scheme. In our case, each parameter can be updated in a Gibbs sampling step, except the coefficients from the slab component, for which we use adaptive rejection sampling as implemented in the `R` package `armspp`. Details are in the supplementary materials.

### 4.5 Simulations

In order to build realistic simulations, in the setting without covariates, we based on our simulated datasets on real data from the Pan-Cancer Analysis of Whole Genomes (PCAWG) study [icg, 2020]. In particular, we ran standard, single-study NMF approaches on 10 diverse cancer types with a range of sample sizes to identify signatures that are likely to be present in each. We then used the published signatures from COSMIC representing the closest matches to these estimates as ground-truth, and estimated exposures from the real data conditional on those signatures. With these signatures and exposures, we finally generated data under a Poisson likelihood according to our model.

In the 3-study simulation, we generated data in this way using the PCAWG data for lung adenocarcinoma, soft tissue leiomyosarcoma, and soft tissue liposarcoma, which resulted in data consisting of 38, 15, and 19 samples with 11, 8, and 5 ground-truth signatures respectively. Across these three simulated studies, five signatures were common, five were study-specific, and the remaining two were partially shared. In total, we generated 10 instances of this 3-study simulation with identical ground-truth parameters each time.

The 10-study simulation was built using these same three PCAWG datasets, as well as data for glioblastoma (41 samples and 5 ground-truth signatures), medullablastoma (146 samples and 6 ground-truth signatures), oligodendroglioma (18 samples and 4 ground-truth signatures), pilocytic astrocytoma (89 samples and 5 ground-truth signatures), renal cell carcinoma (144 samples and 8 ground-truth signatures), chromophobe renal cell carcinoma (45 samples and 8 ground-truth signatures), and squamous cell cancer (48 samples and 6 ground-truth signatures). Two signatures are common to all of the studies, nine are study-specific, and the rest, which comprises the majority, are partially shared with many various configurations. We again generated a total of 10 instances of this 10-study simulation.

In the setting with covariates, again to build realistic simulations, we based our simulated datasets on our colorectal cancer application, in which we have three studies with 16, 62, and 328 samples respectively. We applied single-study NMF approaches to these data to identify signatures with close matches to those in COSMIC, leading us to include a total of four signatures: two that are partially shared and two that are study-specific. We also looked for associations between covariates and exposures to these signatures, which motivated modeling effects from gender and age. Specifically, we generated gender indicators from a Bernoulli(0.5) distribution and age from a Normal(0, 1) distribution (representing z-scored age). We then generated exposures under our covariate model. For the study with 16 samples, male gender was positively associated (*δ* = 1.5) with two signatures, and age was negatively associated (*δ* = −1.5) with the same two signatures. For the other two studies, there were no associations for either covariate with any signature.

Further details are in the supplementary materials.

#### 4.5.1 Evaluation and Comparison to Existing Methods

In the setting without covariates, we evaluated the performance of our approach through three metrics: cosine similarity for the signature estimates, and sensitivity and precision for the sharing pattern. In particular, we computed the cosine similarity between each estimated signature and those in the COSMIC catalogue. We labeled each signature by the name of its closest match in COSMIC, as long as the cosine similarity was over 0.70. If a signature estimate did not have any cosine similarities over that threshold, we considered that signature “unlabeled.” We then used these labels to compute sensitivity and precision. Sensitivity was reported as the total number of labeled signatures found in each study that should have been there according to ground truth, divided by the total number of, not necessarily unique, signatures that should be present across all studies. Precision was found using the same numerator, but divided by the total number of signatures, labeled and unlabeled, found in each study. Note that if multiple estimated signatures were given the same label, at most one would be considered correct and the rest were considered incorrect.

We ran these same simulations under existing approaches and computed the same metrics on the results as a comparison. For both HDP [Roberts, 2018] and the corresponding semi-supervised approach, which we call HDP-frozen, we used the default prior and sampler specifications provided in the online tutorial. Because it is not straightforward to produce posterior summaries, Roberts [2018] suggests a post-hoc clustering algorithm on the MCMC samples to produce consensus signatures and corresponding estimates. Since our results are reported using direct posterior output, we chose not to apply this post-processing algorithm to the HDP output and to instead report the modal sensitivity and precision from the MCMC samples.

For the single-study approaches, signeR [Rosales et al., 2017] and SigProfiler [Alexandrov et al., 2020], we ran each approach on one study at a time within each simulation. These methods were run with default specifications, and we used their reported point estimates of the signatures when computing our evaluation metrics.

## 4.6 Code Availability

R code implementing our methods can be found at www.github.com/igrabski/multi_study_nmf.

